# Abnormal Neurite Density and Orientation Dispersion in Frontal Lobe Link to Elevated Hyperactive/Impulsive Behaviors in Young Adults with Traumatic Brain Injury

**DOI:** 10.1101/2021.08.18.456850

**Authors:** Meng Cao, Yuyang Luo, Ziyan Wu, Kai Wu, Xiaobo Li

## Abstract

Traumatic brain injury is a major public health concern. A significant proportion of individuals experience post-traumatic brain injury behavioral impairments, especially in attention and inhibitory control domains. Traditional diffusion-weighted MRI techniques, such as diffusion tensor imaging, have provided tools to assess white matter structural disruptions reflecting the long-term brain tissue alterations associated with traumatic brain injury. The recently developed neurite orientation dispersion and density imaging is a more advanced diffusion-weighted MRI modality, which provides more refined characterization of brain tissue microstructures by assessing the neurite orientation dispersion and neurite density properties. In this study, we investigated the morphometrical and microstructural alterations at chronic brain injury stage and their relationships with the functional outcomes. Neurite orientation dispersion and density imaging data from 44 young adults with chronic traumatic brain injury (ranging from 18 - 27 years of age; 23 males/21 females) who had no prior-traumatic brain injury history of attention deficits and/or hyperactivity and 45 group-matched normal controls (23 males /22 females) were collected. Maps of fractional anisotropy, neurite orientation dispersion index, and neurite density index were calculated. Vertex-wise and voxel-wise analyses were conducted for gray matter and white matter, respectively. Post-hoc region of interest-based analyses were also performed. Compared to the controls, the group of traumatic brain injury showed significantly increased orientation dispersion index in various gray matter regions and significantly decreased orientation dispersion index in several white matter regions. Brain-behavioral association analyses indicated that the reduced neurite density index of left precentral gyrus and the reduced orientation dispersion index of left superior longitudinal fasciculus were significantly associated with elevated hyperactive/impulsive symptoms in the patients with traumatic brain injury. These findings suggest that traumatic brain injury-induced chronic neurite orientation dispersion alterations of left superior longitudinal fasciculus and left precentral may significantly contribute to post-traumatic brain injury hyperactive/impulsive behaviors in young adults with traumatic brain injury.

## Introduction

Traumatic brain injury (TBI) is a major public health concern, with about 2.8 million documented cases occurring each year in the United States,^1^ approximately 10 percent of which are due to sports and recreational activities.^2^ Although the acute symptoms of TBI may be transient and temperate, evidence is emerging that a significant proportion of patients experience long-lasting post-TBI cognitive, emotional, behavioral, sensory, and motoric changes.^3^ Among those post-TBI cognitive and behavioral disturbances, attention deficits are the most commonly reported.^4^ with the neuropathological natures remaining unclear.

The long-term post-TBI neural and psychophysiological anomalies have been suggested to relate to traumatic axonal injury that persists in a chronic stage.^5^ Loss of white matter (WM) integrity has been widely considered to play a major role in the clinical phenomenology of TBI.^6^ The pathology of diffuse axonal injury has been widely investigated in diffusion MRI (dMRI)-based approaches, especially diffusion tensor imaging (DTI) that is more sensitive than conventional T1- or T2-weighted imaging in detecting axonal injury.^7, 8^ Among the DTI metrics, fractional anisotropy (FA) has been one of the most commonly reported measures in existing studies in TBI, with findings highly inconsistent in terms of involved brain regions as well as the trends (increase or reduction) of FA abnormalities. In particular, a number of existing studies indicated reduced FA in commonly affected WM tracts including corticospinal tract, ^9^ sagittal stratum,^10^ and superior longitudinal fasciculus (SLF)^11^; corona radiata,^12^ uncinated fasciculus,^10, 13^ corpus callosum,^12^ inferior longitudinal fasciculus,^13^ and cingulum bundle^13^; inferior fronto-occipital fasciculus (IFOF)^14^ among patients with chronic TBI relative to controls. While increased FA in corpus callosum,^15^ internal capsule,^16^ corticospinal tracts,^15^ corona radiate^16^ were also reported in subjects with a history of TBI as compared with matched controls.

The FA value derived from DTI can be influenced by changes in axial diffusivity (AD), radial diffusivity (RD) within the brain tissues, or both, making FA sensitive to many types of brain pathology, while incapable for identifying the specific nature of the underlying pathology.^17^ In addition, although DTI can describe water diffusion behavior at modest b-values (e.g., 1000 s/mm^2^), many studies have demonstrated that simple Gaussian diffusion models do not sufficiently describe water diffusion in complex tissues, such as crossing fibers and mixtures of diffusion compartments. Thus, conventional DTI approaches lack the ability to distinguish the spatial organization of neurites. Furthermore, multiple mechanisms, including changes in cell morphology and packing density, or changes in WM fiber orientation, would contribute to the alterations of FA. Recently, a more advanced dMRI modality, the neurite orientation dispersion and density imaging (NODDI), has been developed.^18^ This innovative technique acquires imaging data at multiple levels of diffusion weight, with each sampling many different spatial orientations at high angular resolution. The main parameters derived from NODDI are neurite density index (NDI), and neurite orientation dispersion index (ODI). The intracellular component of NODDI is designed to be representative of both axons and dendrites, thus providing an improved description of gray matter (GM) microstructure. In addition, NODDI enables differentiation of changes to tissue density and tissue ODI (both expressed by FA changes when using DTI), ultimately offering superior sensitivity to micro-level tissue changes. NODDI has shown significant promise in the evaluation of stroke,^19^ Alzheimer’s disease,^20^ and other patient populations.^21^ Recently, Churchill et al. conducted a NODDI study in adult athletes with and without TBI and reported significantly increased NDI and decreased ODI in corpus callosum and internal capsule in the TBI group.^22^ Besides, another recent study found that compared to conventional DTI, the NODDI parameters were more sensitive imaging biomarkers underlying WM microstructural pathology for relatively short-term (2 weeks and 6 months after TBI) consequences post mild TBI.^23^ However, the relationships between NODDI measures and the common chronic (> 6 months) post-TBI behavioral alterations have not been investigated.

In this study, we proposed to utilize the refined measures provided by NODDI to investigate the neurobiological mechanisms that may underlie TBI-related attention deficits in young adults with chronic TBI. Based on results of our previous functional near-infrared spectroscopy (fNIRS) studies that showed functional alterations in frontal areas during sustained attention processing in adults with TBI,^24,25^ we hypothesized that neural morphometrical and microstructural abnormalities in frontal areas and the major WM fibers connecting frontal and other brain regions may exist, and significantly contribute to post-TBI attention-related behavioral alterations.

## Materials and Methods

### Participants

A total of 89 young adults (ranging from 18 - 27 years of age), including 44 patients with TBI (23 males/21 females) and 45 group-matched normal controls (NCs; 23 males /22 females), were initially involved in this study. Participants in the TBI group were recruited from sports teams at New Jersey Institute of Technology (NJIT) and Rutgers University (RU). The NCs were recruited through on-campus flyers at NJIT and RU. The study received Institutional Reviewed Board Approval at both NJIT and RU. Written informed consents were provided by all participants. Within these 89 subjects, 27 were involved in our previous fNIRS study.^24^

The subjects with TBI had a history of one or multiple sports- or recreational activity-related non-penetrating TBIs, with the most recent onset of TBI clinically confirmed at least 6 months prior to the study visit; had no head injury which ever caused overt focal brain damage; had no history of diagnosis with any sub-presentation of attention-deficit/hyperactivity disorder (ADHD) prior to the first onset TBI. The group of NCs included young adults with no history of head injury; had no history of diagnosis with ADHD; and had no severe inattentive and/or hyperactive/impulsive behaviors measured using the Conner’s Adult ADHD Self-Reporting Rating Scales (CAARS, T-score < 60 for both inattention and hyperactivity/impulsivity subscales) during the study visits. Subjects in both groups were native or fluent speakers of English and strongly right-handed based on the Edinburgh Handedness Inventory. The study excluded subjects who had a history or current diagnosis of any neurological disorders (such as epilepsy); severe psychiatric disorders (including Schizophrenia, Autism Spectrum Disorders, Major Depression, Anxiety, etc.); received treatment with any stimulant or non-stimulant psychotropic medication within the month prior to testing; or had contraindications to MRI scanning. Demographic and clinical/behavioral information of the study cohort was included in **Table 1**.

**Table 1:** Demographic and clinical characteristics in groups of normal controls and traumatic brain injury.

|  | **NC (N=40)**<br>**Mean (SD)** | **TBI (N=43)**<br>**Mean (SD)** | ***p*** |
| --- | --- | --- | --- |
| **Age (year)** | 21.40 (2.31) | 21.02 (2.04) | *0.432* |
| **Education Level (year)** | 14.70 (1.73) | 14.23 (1.51) | *0.192* |
| **Mother's Education Level (year)** | 15.36 (2.21) | 15.76 (2.62) | *0.467* |
| **Father's Education Level (year)** | 15.92 (2.99) | 15.81 (2.36) | *0.849* |
| **Conners' Adult ADHD Rating Scale (T score)** |  |  |  |
| Inattentive | 45.53 (6.43) | 57.67 (15.28) | *<0.001* |
| Hyperactive/Impulsive | 43.25 (5.65) | 52.26 (13.72) | *<0.001* |
| ADHD Total | 44.03 (6.14) | 56.67 (16.36) | *<0.001* |
| **Conners' Adult ADHD Rating Scale (raw score)** |  |  |  |
| Inattentive | 4.50 (2.78) | 9.53 (6.40) | *<0.001* |
| Hyperactive/Impulsive | 5.38 (2.59) | 9.02 (5.39) | *<0.001* |
| ADHD Total | 9.88 (4.69) | 18.44 (10.65) | *<0.001* |
|  | **N (%)** | **N (%)** | ***p*** |
| **Male** | 22 (55.0) | 23 (53.5) | *0.89* |
| **Right-handed** | 40 (100.0) | 43 (100.0) | *1.00* |
| **Race** |  |  | *0.127* |
| *Caucasian* | 13 (32.5) | 22 (51.2) |  |
| *African American* | 3 (7.5) | 7 (16.3) |  |
| *Hispanic/Latino* | 2 (5.0) | 2 (4.7) |  |
| *Asian* | 17 (42.5) | 8 (18.6) |  |
| *More than one race* | 5 (12.5) | 4 (9.3) |  |

TBI: group of traumatic brain injury; NC: group of normal controls; SD: standard deviation

Four subjects were excluded from group-level analysis due to the heavy head motion (with either the mean relative volume-to-volume displacement, maximum rotation, or maximum translation > 2.5mm).

### Neuroimaging Data Acquisition Protocol

High-resolution T1-weighted structural MRI (0.9 mm^3^ isotropic T1w and T2w images) and dMRI data were acquired from a 3.0T Siemens Trio imaging system (Siemens, Erlangen, Germany). The three-dimensional dMRI images were acquired using a three-shell protocol with an echo planar imaging pulse sequence: TR/TE = 7700/103 ms, voxel size = 2.0 × 2.0 × 2.5 mm^3^, number of slices = 55, FOV = 220 × 220 × 138 mm^3^. The three shells of dMRI were consisted of 64 diffusion-weighting directions acquired at b = 300 s/mm^2^, b = 700 s/mm^2^, and b = 2000 s/mm^2^, along with six b0 scans. Data of the b = 700 s/mm^2^ shell were used for both DTI and NODDI analyses.

### Individual-level Imaging Data Processing

T1-weighted MRI data preprocessing were conducted using the Human Connectome Project (HCP) pipeline^26^ and FreeSurfer v6.0.0.^27^ After correction of gradient nonlinearity and field intensity, individual white and pial surfaces for surface-based analysis in GM were generated in the native spaces. Then each individual’s pial surface was non-linearly registered to the group-averaged pial surface using multimodal surface matching (MSM) algorithm.^28^

Each dMRI data was first preprocessed using the FMRIB Software Library (FSL; www.fmrib.ox.ac.uk/fsl) for corrections of gradient distortion, B0 distortion and eddy current distortion.^29^ The transformation matrices from local diffusion space to the structural space were calculated by registering a b0 volume to the T1-weighted image using Freesurfer’s *BBRegister*.^30^ Voxel-wise NODDI analyses were then conducted using the open-source MATLAB toolbox (http://mig.cs.ucl.ac.uk/index.php?n=Tutorial.NODDImatlab) with the “WatsonSHStickTortIsoV_B0” parameterization to distinguish three microstructural subcomponents: intracellular (neuritic), extracellular, and cerebrospinal fluid (CSF) compartments. The NODDI MATLAB toolbox was utilized to generate three parameter maps: 1) fraction of CSF (f_CSF_), which indexes the volume fraction of Gaussian isotropic diffusion (free fluid) within each voxel; 2) NDI, which indicates the fraction of tissue water restricted within neuritis (axons and dendrites) in the non-CSF compartment. The intracellular signal was modelled as a Watson distribution over cylinders of zero radius, which has a mean orientation vector *µ* and a concentration parameter *κ* ∈ (0, ∞) indicating how much the distribution tends to spread out around *µ*; and 3) ODI, transformed from the concentration parameter, which characterizes spatial configuration of neurites and ranges from 0 (completely parallel neurites) to 1 (completely random neurite orientation). In addition, the b = 700 s/mm^2^ data series were analyzed to extract voxel-wise conventional DTI measure, FA, by using FSL’s *dtifit*. The individual-level data processing steps were shown in **Fig. 1A**.

**Figure 1.**
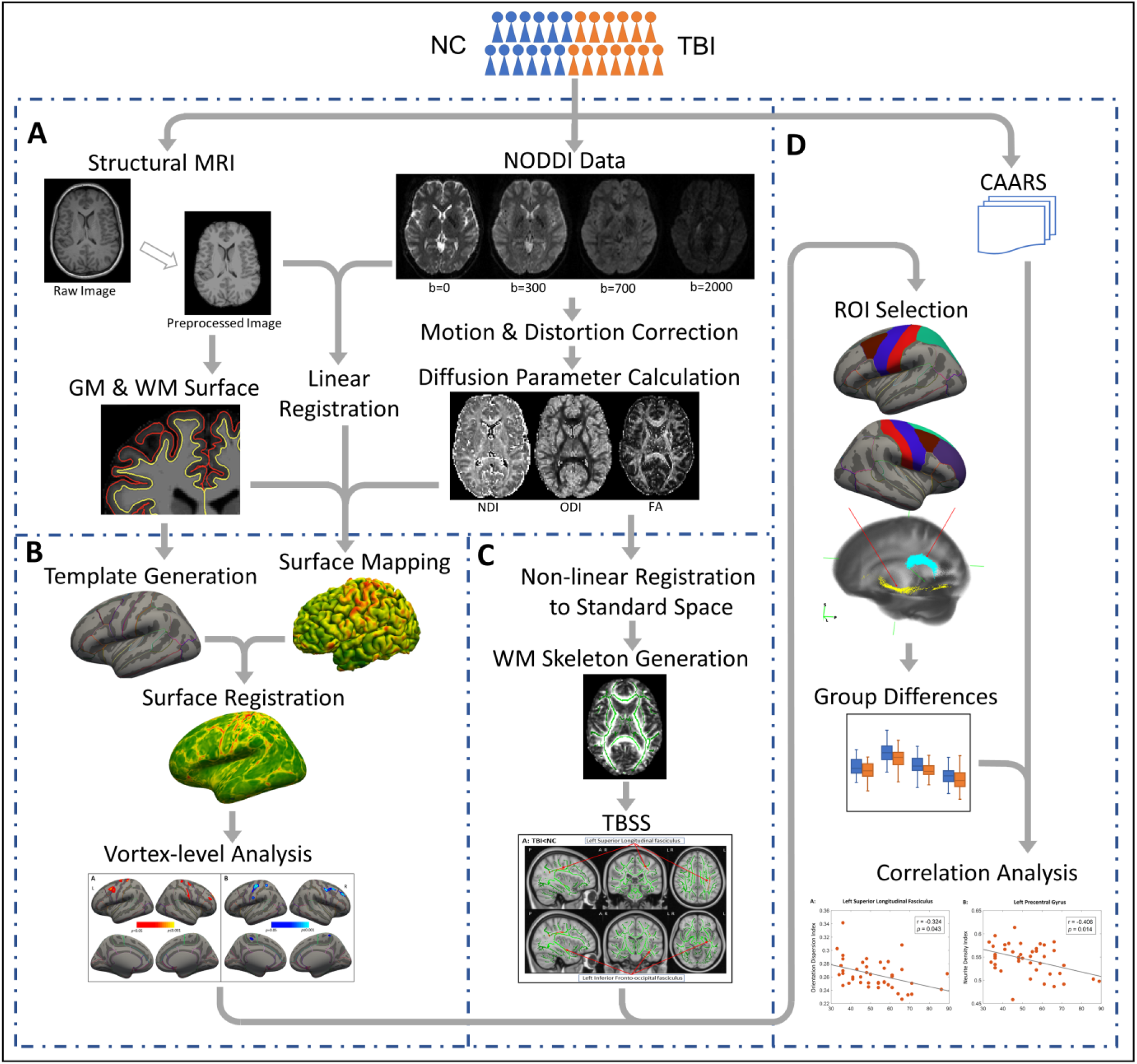
Overview of the study methods. **(A)** Individual-level data processing. **(B)** Surface-based analysis of GM. **(C)** Voxel-based analysis of WM. **(D)** Group-level statistical analysis. NC: normal controls; TBI: traumatic brain injury; NODDI: neurite orientation dispersion and density imaging; GM: gray matter; WM: white matter; TBSS: tract-based spatial statistics; CAARS: Conner’s adult attention-deficit/hyperactivity disorder self-reporting rating scales; ROI: region of interest.

### Surface-Based Analysis in GM

The surface-based analysis in GM was conducted using the HCP’s connectome workbench and Freesurfer. First, the NDI, ODI, and FA maps of each individual were linearly transformed into native structural space. A ribbon mapping method, using WM surface as inner surface and pial surface as outer surface, was applied to sample the diffusion measures onto each individual’s surface at local space. This method was to construct a polyhedron for each vertex using the vertex’s neighbors, and to calculate the weight of each vertex based on the overlapping volumes with nearby voxels. To generate the group difference map, the surface maps of NDI, ODI, and FA were subsequently resampled onto the group-averaged surfaces (**Fig. 1B**) based on MSM surface registration.^28^

The surface-based group comparison between TBI and NCs were performed using the Freesurfer’s GLM analysis. Age, sex, and parent education level were used as covariates. A cluster-wise correction method, with a cluster of 25 mm^2^ or greater and p value < 0.05, were performed, as the thresholding method for the vertex-wise group difference map. The anatomical regions with significant group differences on the GM surface map were identified using Desikan-Killiany atlas.

### Tract-Based Spatial Statistics (TBSS) Analysis in WM

TBSS of the NDI, ODI and FA maps were carried out in FSL (version 6.0.1). Firstly, FMRIB58_FA in the Montreal Neurological Institute (MNI) common space was used as the target image for nonlinear registration of all subjects’ FA maps, using the FMRIB Nonlinear Registration Tool (FNIRT; http://fsl.fmrib.ox.ac.uk/fsl/fslwiki/FNIRT/). The transformed FA images in the MNI space were then averaged and skeletonized, to generate the major WM skeleton that would be implemented to all the subjects (**Fig. 1C**). The threshold of FA ≥ 0.2 was implemented to the average FA map to ensure the WM skeleton to include major WM tracts while exclude peripheral tracts and GM. Each participant’s aligned FA map was then projected onto this skeleton by assigning to each voxel the maximum FA in a line perpendicular to the local skeleton. The NDI, ODI, and isotropic compartment were projected onto the mean FA skeleton after applying the warping registration field of each subject to the standard MNI space.

Comparisons between the groups of TBI and NCs were performed by voxel-wise statistics of the skeletonized maps of NDI, ODI and FA, using nonparametric statistical thresholding approach (FSL Randomise permutation algorithm; https://fsl.fmrib.ox.ac.uk/fsl/fslwiki/Randomise). The thresholded mean FA skeleton was used as a mask. Two thousand permutations and statistical inference using threshold-free cluster enhancement (TFCE) were performed, with p values < 0.05 after family-wise error (FWE) correction for multiple comparisons. Age, sex, and parent education level were used as covariates. The anatomic locations of regions with significant group differences on the WM skeleton were identified from the Johns Hopkins University WM labels atlas.

### Post-hoc Region of Interest (ROI) Analysis in GM and WM

On the base of the intermediate voxel-based results from the surface-based analysis in GM and TBSS in WM, post-hoc region of interest (ROI)-based analyses were further performed, as shown in **Fig. 1D**. A total of 13 GM and WM brain regions that showed significant between-group differences in the voxel-based analyses, including 11 GM ROIs (right rostral middle frontal gyrus, right superior frontal gyrus, left superior parietal lobule, bilateral caudal middle frontal gyrus, bilateral precentral gyrus, bilateral postcentral gyrus, and bilateral paracentral lobules) and two WM ROIs (left IFOF and left SLF), were involved in ROI-based analysis. The GM and WM ROIs were defined based on the Desikan-Killiany atlas and the Johns Hopkins University WM probabilistic tractography atlas, respectively. Mean ODI and NDI values within each GM and WM ROI was extracted for each subject (no between-group differences in FA were found during the voxel-based analyses). One-way analysis of covariates (ANCOVA) was conducted to identify the between group differences, with age, sex, and parent education level as covariates. Bonferroni correction was applied to control multiple comparisons at significance level of 0.05.

### Brain-Behavior Correlation Analysis

Partial correlation was conducted between the ROI-based brain imaging measures that showed significant between-group differences and the behavioral measures for clinical symptoms in inattentiveness and hyperactivity/impulsivity (measured using the T-scores of the CAARS inattentive and hyperactive/impulsive subscales) Again, Bonferroni correction was applied to control multiple comparisons at significance level of 0.05.

## Results

### Demographic, Clinical and Behavioral Measures

The demographic and clinical information of both TBI and NC groups were summarized in **Table 1**. Demographic measures did not show significant between-group difference. Compared to the NCs, subjects with TBI showed significantly more clinical symptoms in inattentiveness and hyperactivity/impulsivity measured using the T- and raw-score of the CAARS inattentive and hyperactive/impulsive sub-scales.

### Group Differences in the Intermediate Voxel-based Analyses

Group-level results of the surface-based analysis in GM showed significantly decreased NDI in vertex clusters from bilateral precentral gyrus, bilateral postcentral gyrus, left caudal middle frontal, right superior frontal gyrus, and right rostral middle frontal gyrus in patients, relative to controls (**Fig. 2A**). In addition, TBI group demonstrated significantly increased ODI in clusters of bilateral precentral gyrus, bilateral paracentral lobules, left precentral gyrus, left superior parietal lobule, and right middle frontal gyrus, when compared to NCs (**Fig. 2B**). All these regions showed *p*-value < 0.05 with a cluster greater than 25 mm^2^.

**Figure 2.**
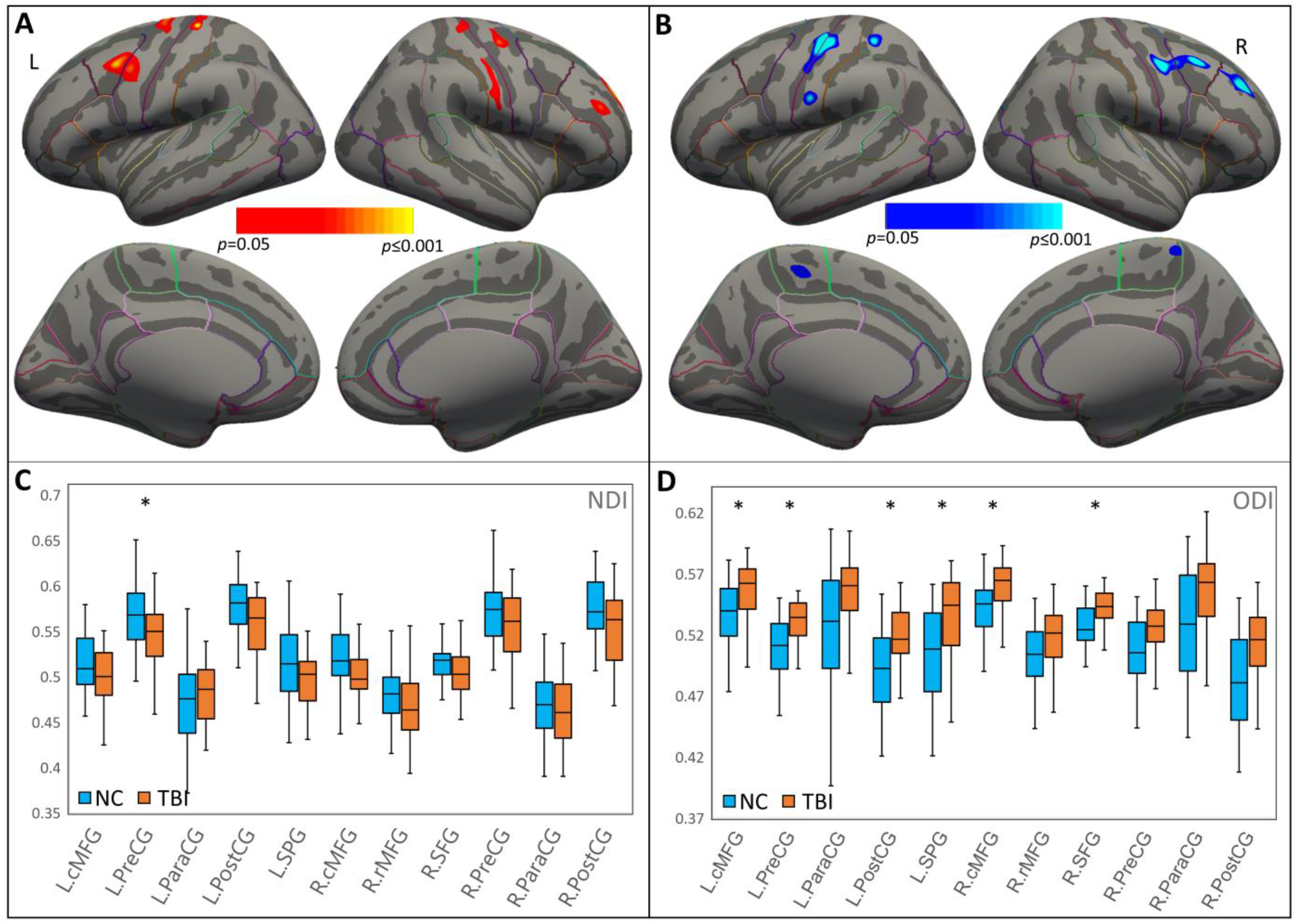
Surface-based analysis of neurite properties in gray matter. **(A)** Surface-based gray matter analysis showing clusters that have higher NDI in NC than TBI. **(B)** Surface-based gray matter analysis showing clusters that have lower ODI in NC than TBI. **(C)** Group comparison of NDI in selected brain regions. Regions that showed significant group difference after Bonferroni correction were marked with asterisk (*). **(D)** Group comparison of ODI in selected brain regions. Regions that showed significant group difference after Bonferroni correction were marked with asterisk (*). NDI: neurite density index; ODI: orientation dispersion index; TBI: traumatic brain injury; NC: normal controls; L: Left; R: Right; cMFG: caudal middle frontal gyrus; PreCG: precentral gyrus; ParaCG: paracentral gyrus; PostCG: postcentral gyrus; SPG: superior parietal gyrus; rMFG: rostral middle frontal gyrus; SFG: superior frontal gyrus.

Group-level results of TBSS in WM showed significantly decreased ODI in the left SLF (*p* < 0.05, TFCE corrected) and left IFOF (*p* < 0.05, TFCE corrected) in patients with TBI relative to NCs (**Fig. 3A**).

**Figure 3.**
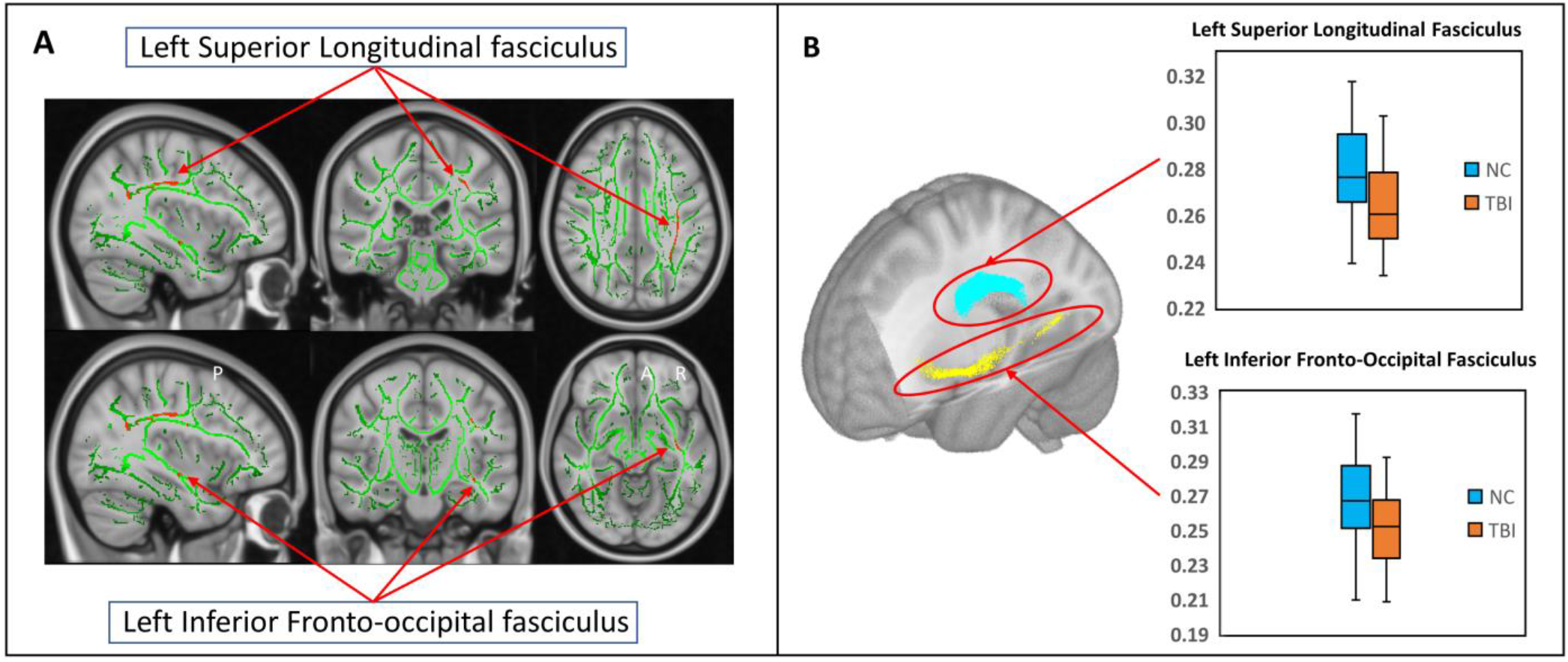
Voxel-level and region of interest analyis of white matter. **(A)** Tract-based spatial statistics results showing significant between-group differences of voxel-wise ODI for groups of TBI and normal controls. **(B)** Region of interest analysis results showed significant between group differences in the groups of NC and TBI. Top panel presents the results in left superior longitudinal fasciculus; bottom panel presents the results in left inferior fronto-occipital fasciculus. TBI: traumatic brain injury; NC: normal controls; ODI: orientation dispersion index.

### Group Differences of ROI-based Imaging Measures

Compared to controls, the TBI group showed significantly abnormal neurite orientational integrity in bilateral frontal and parietal GM areas, represented by greatly increased ODI in bilateral middle frontal gyri, postcentral, precentral, and superior parietal gyri in the left hemisphere, as well as the superior frontal gyrus in the right hemisphere. Relative to controls, the adults with TBI also showed significantly decreased GM neurite density left precentral gyrus. The ROI-based analyses in WM showed significantly reduced ODI in left IFOF and SLF in the group of TBI. All the parameters of these results are detailed in **Table 2** and graphically depicted in **Fig. 2C&D** and **Fig. 3B**, with all the *p* values corrected using Bonferroni method.

**Table 2:** Anatomical regions that showed significant between-group differences of the neurite morphometry.

| Regions | Measures | NC (N=40)<br>Mean (SD) | TBI (N=43)<br>Mean (SD) | *F* | *p*# |
| --- | --- | --- | --- | --- | --- |
| **Gray Matter ROIs** |  |  |  |  |  |
| Left Caudal Middle Frontal | ODI | 0.537 (0.024) | 0.556 (0.022) | 18.17 | <0.001 |
| Left Postcentral Gyrus | ODI | 0.493 (0.035) | 0.517 (0.027) | 11.296 | 0.014 |
| Left Precentral | ODI | 0.509 (0.024) | 0.529 (0.022) | 19.743 | <0.001 |
|  | NDI | 0.568 (0.036) | 0.545 (0.033) | 9.903 | 0.026 |
| Left Superior Parietals | ODI | 0.503 (0.039) | 0.535 (0.035) | 12.813 | 0.007 |
| Right Caudal Middle Frontal | ODI | 0.542 (0.023) | 0.559 (0.021) | 15.024 | 0.003 |
| Right Superior Frontal | ODI | 0.529 (0.017) | 0.542 (0.015) | 16.512 | 0.001 |
| **White Matter ROIs** |  |  |  |  |  |
| Left Inferior Fronto-occipital Fasciculus | ODI | 0.266 (0.028) | 0.251 (0.021) | 8.115 | 0.012 |
| Left Superior Longitudinal Fasciculus | ODI | 0.276 (0.022) | 0.264 (0.022) | 6.039 | 0.032 |

ROI: region-of-interest; TBI: group of traumatic brain injury; NC: group of normal controls; SD: standard deviation; ODI: orientation dispersion index; NDI: neurite density index; *p*#: *p* value after Bonferroni correction.

### Brain-Behavior Correlations

In the group of TBI, reduced NDI of the left precentral gyrus and reduced ODI of the left SLF were both significantly correlated with elevated hyperactive/impulsive symptoms (**Fig. 4**). We did not find significant brain-behavior correlations in the group of NC.

**Figure 4.**
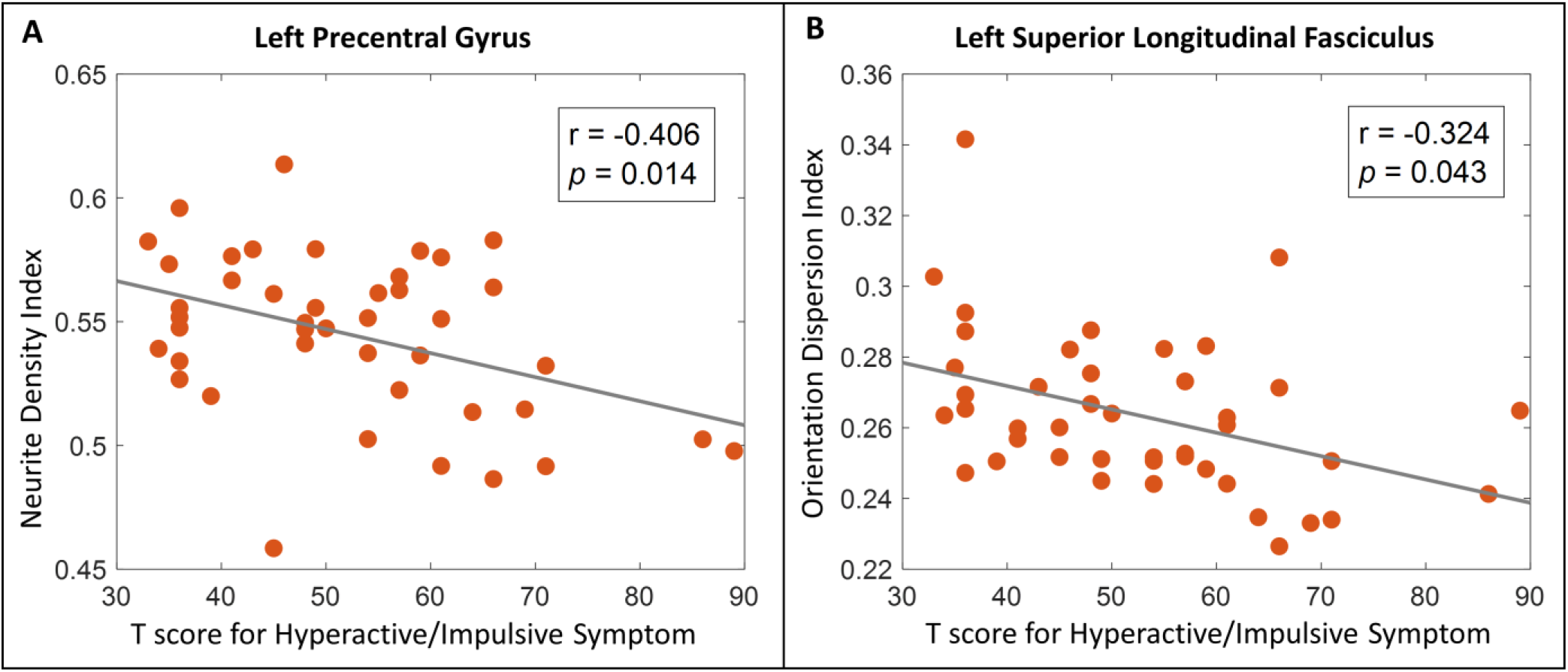
Brain-behavior correlation analysis results. **(A)** Greater orientation dispersion index of left superior longitudinal fasciculus was significantly correlated with reduced hyperactive/impulsive symptom severity T-score in the group TBI. **(B)** Increased neurite density index of left precentral gyrus was significantly correlated with reduced hyperactive/impulsive symptom severity T-score in the group TBI. TBI: traumatic brain injury.

## Discussion

The present study investigated the neurite morphometry differences, represented by the NDI and ODI in both GM and WM areas, between young adults with TBI and group-matched controls. The results in surface-based GM analysis reported significantly altered neurite morphometry in bilateral frontal and parietal areas, with the left frontal morphometrical abnormalities to be dominated in the subjects with TBI. In the literature of TBI-related studies, structural and functional abnormalities associated with frontal GM areas have been consistently reported in adults and children with TBI. For instance, relative to controls, TBI groups showed significantly decreased thickness^32-34^ and significantly increased activation during working memory task^35, 36^ in frontal lobe. In addition, cortical thinning^37, 38^ and hyperactivation^35, 39^ in parietal lobe were also frequently observed in adults and children with TBI. Along with existing findings, our results suggest frontal and parietal regions are highly susceptible to the TBI-induced axonal and neuronal damages and NODDI have the high sensitivity in detecting the microstructure alterations.

Our investigations in WM demonstrated that compared to matched controls, the subjects with TBI had significantly decreased ODI in the left SLF and left IFOF. The SLF and IFOF are two key components of the long-association fronto-parietal pathways, which interconnect cortical and subcortical areas to subserve the visuospatial attention and higher-order cognitive processes.^40, 41^ Studies have suggested that SLF and IFOF are critical WM structures involving in the visuospatial orienting and executive components of attention processing,^42^ and brain regions associated with the orienting and executive components of visuospatial attention are most vulnerable to neural damages resulting from mild TBI.^43^ Indeed, substantial previous DTI studies in subjects with TBI have reported structural abnormalities of SLF and IFOF. For instance, significantly increased FA and decreased mean diffusivity (MD) in SLF and/or IFOF have been frequently reported in subjects with a history of sports-related concussion.^15, 16, 22^ While other studies in subjects with TBI have reported decreased FA and/or increased MD in left SLF and/or IFOF, and their linkages with prolonged post-TBI behavioral impairments.^44-46^ The inconsistence of these existing findings may be partially explained by the long-term effects of TBI appear to differ from subacute concussion or chronicity of the injury,^47^ the severity level of TBI,^48^ and the limitation of DTI measures.^49^

Furthermore, the results of the present study implicated that the significantly decreased NDI of left precentral gyrus and decreased ODI of the left SLF both significantly contribute to the elevated hyperactive/impulsive symptoms in individuals with TBI. These findings first time in the field suggest that both the TBI-related reduced GM neurite density in left frontal lobe and the overly aligned axons^18^ in left SLF that anatomically interconnecting frontal and parietal lobes can significantly contribute to TBI-related behavioral impairment in the domain of inhibitory control. The precentral gyrus has been consistently implicated in response inhibition in both primate and human studies.^50-53^ The SLF has also emerged as one of the most reliably identified WM tract underlying inhibitory,^54-57^ and variability in behavioral performances of inhibitory control-related tasks can be partially explained by variability in WM integrity of SLF.^58^ Together with existing studies, our findings suggest that neurite density and orientation dispersion alterations in left frontal lobe, especially the precentral area, are significantly vulnerable in TBI-induced chronic neuronal damages; and these neuronal abnormalities significantly link to post-TBI behavioral impairments especially in the domain of inhibitory control.

## Limitations

Although the current study utilized an innovative and robust dMRI technique to study the neurite morphometry and its relationship with post-TBI behavioral impairments in young adults with TBI and have reported significantly findings, we acknowledge that there are issues to be addressed. First, our cohort consisted of both male and female subjects. Although existing studies indicated that compared to males, females might have poorer consequences after TBI,^59^ our post hoc analyses in subgroups of females with TBI and males with TBI did not finding significant differences in the clinical/behavioral and neuroimaging measures. We acknowledge that the sample sizes of our subgroups were moderate. A future study in a statistically powerful sample is expected to thoroughly investigate the sex-related post-TBI behavioral and anatomical alterations and their interactions. Second, we acknowledge that the diverse injury location of TBI may be associated with different symptoms. Nevertheless, none of the existing studies have identified any patterns of association between a definitive injury location and long-term deficits in a specific domain of cognitive functions.

## Conclusions

By utilizing an innovative dMRI technique, NODDI, the current study investigated the GM and WM neuronal morphometry and its relationship with post-TBI behavioral impairments in young adults with TBI. The results showed that relative to matched controls, the TBI subjects had significantly altered neurite density and orientation dispersion in frontal and parietal GM areas, and the WM tracts connecting these GM areas. We also found strong relationship of the abnormal left precentral GM NDI and SLF WM ODI with increased hyperactive/impulsive behaviors in the group of TBI. Our findings suggest that significantly altered neurite morphometry exists in frontal and parietal GM regions and the WM major tracts that anatomically connect these GM regions, especially in the left hemisphere; and these neurite disruptions significantly contribute to post-TBI attention problems in subjects with TBI. These findings provide valuable insight into the neuropathological markers of chronic TBI, which have the potential to inform biologically targeted prevention and intervention strategies in affected subjects.

## Author Contributions

Dr. Xiaobo Li designed the study. Meng, Yuyang and Ziyan all involved in managing literature search, analyzing the clinical and imaging data, and writing early drafts of the manuscript. Meng and Drs. Li and Wu edited and finalized the manuscript. All authors contributed to and have approved the final manuscript.

